# ADRes: a computational pipeline for detecting molecular markers of Anti-malarial Drug Resistance, from Sanger sequencing data

**DOI:** 10.1101/066183

**Authors:** Setor Amuzu, Anita Ghansah

**Affiliations:** Noguchi Memorial Institute for Medical Research, College of Health Sciences, University of Ghana, P. O. Box LG581, Legon, Ghana

**Keywords:** *Plasmodium falciparum*, antimalarial drug resistance, single nucleotide polymorphisms, molecular markers

## Abstract

**Background:** Malaria control efforts are stifled by the emergence and dispersal of parasite strains resistant to available anti-malarials. Amino acid changes in specific positions of proteins encoded by *Plasmodium falciparum* genes *pfcrt, dhps, dhfr*, and *pfmdr1* are used as molecular markers of resistance to antimalarials such as chloroquine, sulphadoxine-pyrimethamine, as well as artemisinin derivatives. However, a challenge to the detection of single nucleotide polymorphisms (SNPs) in codons responsible for these amino acid changes, in several samples, is the scarcity of automated computational pipelines for molecular biologists to; rapidly analyze ABI (Applied Biosystems) Sanger sequencing data spanning the codons of interest in order to characterize these codons and detect these molecular markers of drug resistance. The pipeline described here is an attempt to address this need.

**Method:** This pipeline is a combination of existing tools, notably SAMtools and Burrows Wheeler Aligner (BWA), as well as custom Python and BASH scripts. It is designed to run on the UNIX shell, a command line interpreter. To characterize the codons associated with anti-malarial drug resistance (ADR) in a particular gene using this pipeline, the following options are required; a path to reference coding sequence of the gene in FASTA format, gene symbol (pfcrt, pfmdr1, dhps or dhfr), and a path to the directory of ABI sequencing trace files for the samples. With these inputs, the pipeline performs base calling and trimming, sequence alignment, and alignment parsing.

**Results:** The output of the pipeline is a CSV (Comma-separated values) file of sample names, codons and their corresponding encoded amino acids. The data generated can be readily analyzed using widely available statistical or spreadsheet software, to determine the frequency of molecular markers of resistance to anti-malarials such as chloroquine, sulphadoxine-pyrimethamine and artemisinin derivatives.

**Conclusions:** ADRes is a quick and effective pipeline for detecting common molecular markers of anti-malarial drug resistance, and could be a useful tool for surveillance. The code, description, and instructions for using this pipeline are publicly available at http://setfelix.github.io/ADRes.

## Background

Malaria is caused by parasitic protozoans of the genus *Plasmodium* and is commonly transmitted by the bites of infected *Anopheles* mosquitoes. Although mortality rates in the WHO Africa region and worldwide have decreased, since the year 2000 [1], malaria remains a global public health concern, with high morbidity and paediatric mortality in endemic areas like sub-Saharan Africa. Every year, malaria kills over 430,000 children in Africa [1]. Malaria control is hampered, in part, by antimalarial drug resistance (ADR). The emergence and spread of resistant strains of *P. falciparum*, the most lethal human malaria pathogen, is a particular cause for concern. The World Health Organization therefore recommends therapeutic efficacy studies (TES) once every two years, comprising *in vivo, ex vivo*, and *in vitro* tests, to guide antimalarial treatment policy in malaria-endemic countries. In addition to TES, antimalarial drug resistance monitoring with molecular markers also informs antimalarial treatment policy [1]. Signature amino acid changes caused by SNPs in specific codons of some *P. falciparum* genes are associated with resistance to specific anti-malarials. These molecular markers provide a useful way for monitoring resistance to antimalarials such as chloroquine, sulphadoxine-pyrimethamine and artemisinin derivates.

Chloroquine is a 4-aminoquinolone drug that was the first-line antimalarial for decades until widespread *P. falciparum* resistance led to its limited use and replacement with artemisinin-based combination therapies (ACTs). Currently, ACTs are the most widely recommended antimalarial for treatment of uncomplicated *P. falciparum* malaria. Chloroquine resistance has been associated with SNPs in the *P. falciparum* chloroquine resistance transporter (*pfcrt*) gene located on chromosome 7. These SNPs result in a modified PfCRT protein that contributes to drug resistance by decreasing the accumulation of 4-aminoquinolones in the food vacuole as a result of increased efflux of the drug from its site of action[2, 3]. Considering that other anti-malarials including the 4-aminoquinolones (such as amodiaquine and desethylamodiaquine) target this organelle, PfCRT and its associated gene mutations are important for the development of resistance. A number of SNPs in *pfcrt* codons 72, 74, 75, 76, 97, 152, have been associated with chloroquine resistance in *P. falciparum* isolates from Southeast Asia, South America and Africa [4, 5]. The mutation at codon 76 resulting in the substitution of threonine for lysine (*pfcrt* K76T) is a key marker of *P. falciparum* chloroquine resistance [5, 6].

Mutations in the *P. falciparum* multidrug-resistance (*pfmdr1*) gene, located on chromosome 5, that are linked to antimalarial drug resistance (ADR) phenotype occur in codons 86, 184, 1034, 1042 and 1246 [7, 8]. This gene, like *pfcrt*, codes for a food vacoule localized protein. The P-glycoprotein homologue-1 (Pgh-1) protein, unlike PfCRT, has been proposed to pump drugs from the cytoplasm towards the food vacuole [9], thereby protecting the parasite from antimalarials with cytoplasmic targets. These anti-malarials include the aminoalcohol-quinolines (AAQs) [10], and artemisinin and its derivatives [11]. Additionally, the substitution of asparagine for tyrosine at position 86 of Pgh-1 has been linked to chloroquine resistance [8]. Infact, mutant *pfmdr1* (not *pfcrt*) has been associated with chloroquine clinical failure in *P. falciparum* malaria in Madagascar [12].

Before the adoption of ACTs, the anti-folate combination of sulfadoxine-pyrimethamine (SP) was popular for treating chloroquine-resistant *P. falciparum* malaria in many African countries [13]. This drug combination functions as an antimalarial by inhibiting the folate synthesis pathway of *P. falciparum* [14]. The enzyme, dihydrofolate reductase (DHFR), which is encoded by the dhfr gene, reduces dihydrofolate to tetrahydrofolate, an important co-factor in nucleic acid and methionine synthesis. Mutations at amino acid positions 50, 51, 59, 108, and 164 of DHFR are associated with pyrimethamine resistance [15, 16]. The substitution of serine for asparagine at position 108 is key for developing pyrimethamine resistance, while the mutations at other positions modulate it [15, 16].

Sulfadoxine resistance is mediated by the accumulation of SNPs in the dihydropteroate synthase (*dhps*) gene which encodes the drug's target (DHPS). Mutations at amino acid positions 436, 437, 540, 581, and 613 in DHPS decrease this enzyme's affinity for sulfadoxine [17–19]. In the absence of these mutations, sulfadoxine is a potent inhibitor of DHPS. Like DHFR, the magnitude of *in vitro* resistance to sulfadoxine is generally associated with the number of amino acid substitutions in DHPS [17, 18]. The substitution of alanine for glycine at position 437, in particular, is considered seminal to sulfadoxine resistance. Triple-mutant DHPS enzymes have been reported to show the highest levels of sulfadoxine resistance in natural parasite populations [17, 18].

Therefore, detecting the SNPs and their corresponding amino acid changes at the respective codons for each of these genes, in several *P. falciparum* isolates, is important for monitoring ADR. Current methods used by our group and others to detect ADR SNPs from sequencing data, typically, involve the use of an alignment viewer to visually count to the position of codons of interest and record their bases, followed by translation of these codons to amino acids. While this manual method produces results, it is not suitable for rapid detection of molecular markers of ADR from hundreds of samples, as it is slow and laborious. To facilitate the rapid detection of molecular markers of ADR in *pfcrt, pfmdr1, dhps*, and *dhfr genes*, we developed an automated computational pipeline to analyze ABI Sanger sequencing data, spanning the relevant codon positions, to produce the codons and their corresponding amino acids for all samples and reference, in a tabular format. This data can be readily analyzed using R, for example, to determine the frequency of molecular markers of resistance to specific antimalarials, including chloroquine, sulphadoxine-pyrimethamine, and artemisinin derivatives.

## Methods

ADRes consists of custom Python and BASH scripts as well as existing tools, namely; Burrows-Wheeler Aligner (BWA) [20], abifpy [21], SAMtools [22], fastq-mcf [23], and sam2fasta.py [24]. These dependencies along with the custom scripts, example data, and instructions for running the pipeline are hosted on GitHub, and are freely available at http://setfelix.github.io/ADRes. ADRes was developed and successfully tested on the UNIX command line. Therefore, a basic tutorial of the UNIX command line [25] is recommended for the uninitiated.

Using ABI Sanger sequencing trace files of regions spanning codons of interest for a given ADR gene, from several *P. falciparum* isolates, this pipeline outputs a CSV file of sample id, codons and their corresponding amino acids. The genes and codons currently supported by ADRes are shown in Table 1.

**Table 1.**
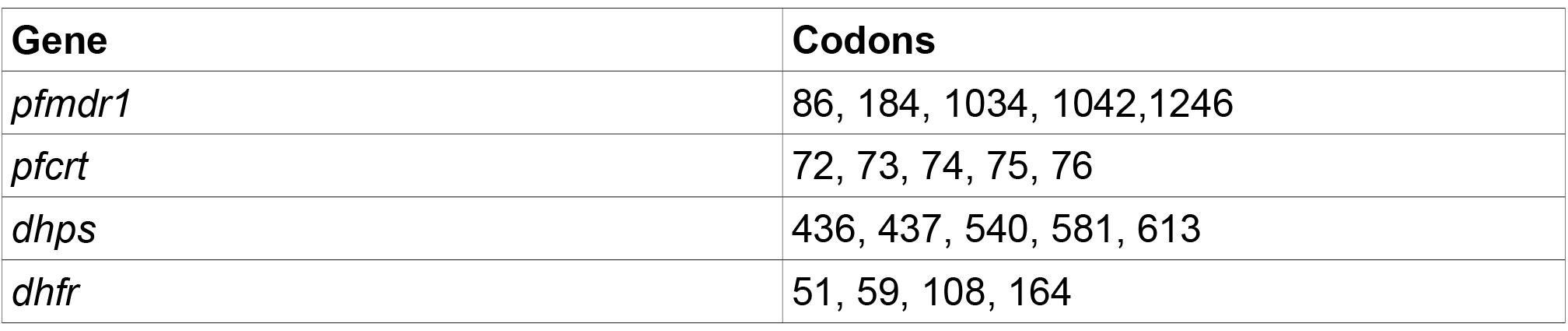
Genes and codons supported by ADRes pipeline

### Installation

1. Download the ADRes package from https://github.com/Setfelix/ADRes/tarball/master and extract it. The package consists of eight python scripts, two BASH scripts, one R script, a README file, a LICENSE file. Additionally, there are three directories (bwa-0.7.12, ea-utils. 1.1.2–806, samtools-1.2) containing the dependencies, and another directory containing example data. Alternatively, the pipeline's GitHub repository can be cloned using this URL: https://github.com/Setfelix/ADRes.git.
2. Optionally, add the ADRes directory to your $PATH variable in bashrc file, for example, so that the pipeline can be run from any directory.

### Usage

The central script for running the pipeline is adres.sh. The following options can be used to run the pipeline:

bash adres.sh <directory_of_ab1_files> <reference_gene_coding_sequence> <gene> [quality_cutoff]

<directory_of_ab1_files>

This option specifies the path to a directory containing ABI Sanger sequencing trace files of a single gene (*dhps, dhfr, pfcrt*, or *pfmdr1*) for several *P. falciparum* isolates. These files have the “.ab1” extension and are DNA sequence chromatograms produced from the Applied Biosystems sequencers, in ABI format [26]. Some example ab1 files are in the 'example ab1' directory, within the 'example_data' directory. This directory contains 50 ab1 trace files obtained from sequencing regions of *pfcrt* spanning codons 72 – 76 from 50 *P. falciparum* isolates.

<reference_gene_coding_sequence>

This option specifies the path to the coding sequence of the respective reference gene, in FASTA format. The reference coding sequence for the *pfcrt* gene [PlasmoDB: PF3D7_0709000] is in the example_ab1 directory. This reference sequence, as well as, those for *dhps, dhfr*, and *pfmdr1* can be downloaded from http://plasmodb.org/plasmo/ [27].

<gene>

One of the following values can be used for this option: pfcrt, dhps, dhfr, pfmdr1. This option instructs the pipeline to use the appropriate parser to generate the CSV output.

[quality_cutoff]

This option specifies the Phred [28] quality threshold causing base removal. Each base call has an associated quality score which estimates the chance that the base call is incorrect. This argument is optional. If no value is given, the default value is 10 (Q10). This corresponds to 1 in 10 chance of incorrect base call. Legal values for this option range from 10 to 60.

### Analysis

An example of a command to analyze *pfcrt* sequencing data, in the 'example_data' directory, using this pipeline is:

bash adres.sh ~/example_data/example_ab1 ~/example_data/pfcrt_pf3D7_cds.fasta pfcrt

The execution of this command can be broken into the following steps:

1. Base calling and quality control – For each ab1 file, the sequence of nucleotide bases and corresponding quality information is retrieved and output to a file in FASTQ [29] format. This is done using abifpy, a python module for reading ABI Sanger sequencing trace files. The resulting FASTQ files are then concatenated into a single file. The fastq-mcf program from the ea-utils package (available at https://code.google.com/p/ea-utils/) is used to remove low quality bases from sample reads using custom or default (Q10) quality threshold.
2. Align trimmed sequences to coding sequence (CDS) of respective reference gene – Prior to alignment, the reference CDS is indexed using bwa index. Alignment is then done using bwa bwasw command, with default settings. This outputs a SAM (Sequence Alignment/Map) [22] file, which can be viewed using Tablet [30] (as shown in figure 1).
3. Alignment filtering and conversion to FASTA format – The SAM alignment is filtered by mapping quality threshold of 10 (MAPQ >= 10), using samtools view. The resulting filtered SAM alignment is then converted to FASTA format using sam2fasta.py. To ensure that sequence lines in the FASTA file are of the same length, the sequence lines for samples are padded with varying number of dots (“….”) depending on their own length. Therefore, the length of the each sequence line in the FASTA file is the same as the length reference sequence. Using this *pfcrt* analysis example, the length of each sequence line in the FASTA file is 1275. The FASTA file contains a single sequence line for every sequence - an example (pfcrt_30_06_15.fasta) can be found in the example_ab1 directory, within the example_data directory. Equal length of sequence lines facilitates counting of codon positions in the next step.
4. Parse FASTA alignment to output codons and corresponding amino acid in CSV format – The FASTA alignment is parsed by the respective python script for the gene under analysis, in this example pfcrt.py is used. The codons, triplet of nucleotides, at the codon positions of interest are retrieved and translated. These values, separated by commas, are then written to a file.

**Figure 1.**
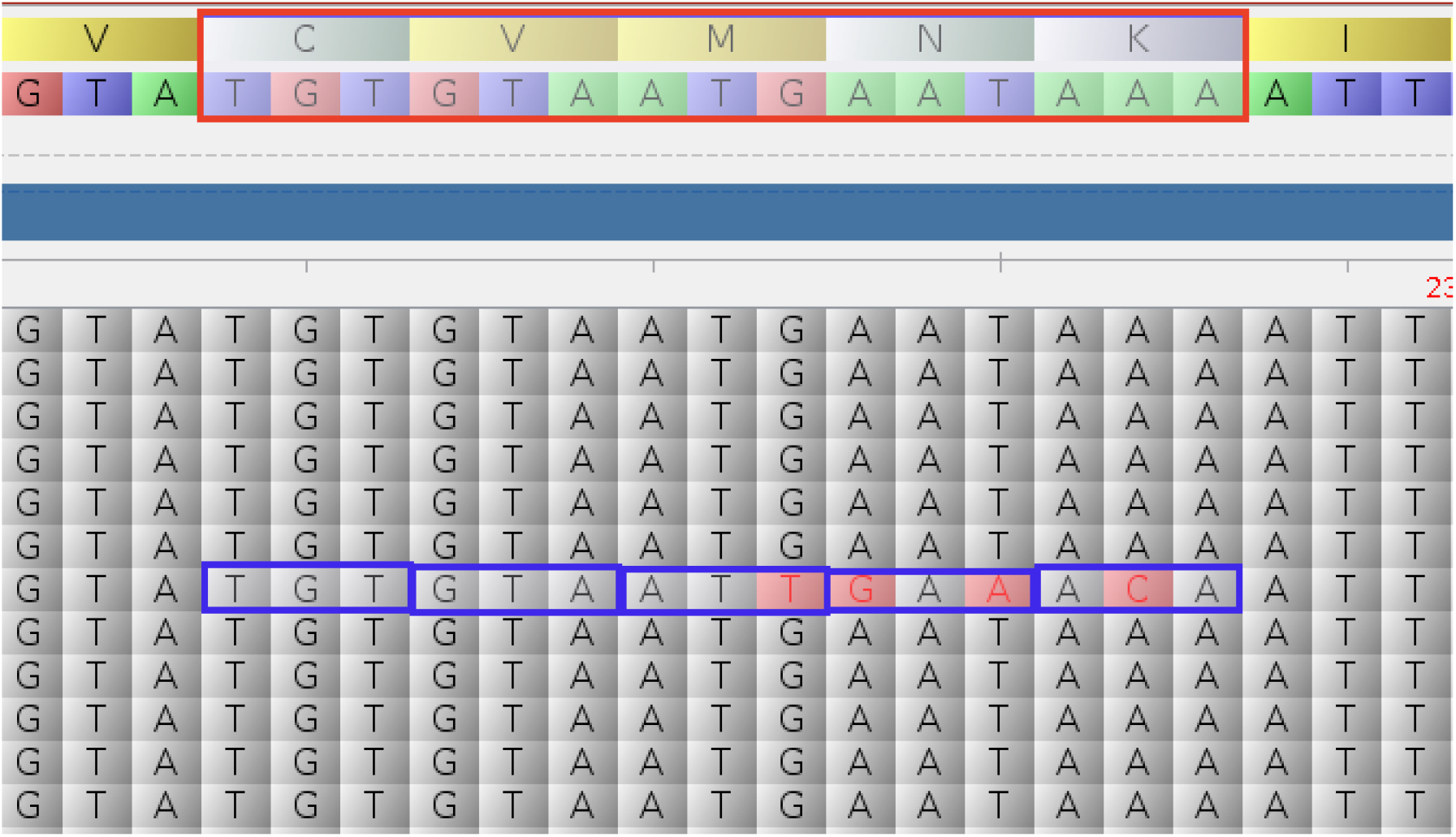
Alignment of codon 72–76 region of *pfcrt* genes from *P. falciparum* isolates to reference CDS of *P. falciparum* 3D7. The reference contains the CVMNK wild-type haplotype (outlined by a red rectangle) which is characteristic of chloroquine-susceptible *P. falciparum.* The blue rectangle outlines a sequence containing SNPs within the codon 72–76 region. Specifically, codons 74, 75, and 76 contain a SNP which altogether produces a CVIET haplotype. Notably, the K76T mutation which is strongly associated with chloroquine-resistance has been detected in this sample – ACA codes for Lysine (T).

The example_ab1 directory contains additional files that are not in ABI format, that is, they do not have the '.ab1' extension. These are output files that were generated after running the pipeline using the example command above. The primary output file is pfcrt_30_06_15.csv. The other output files (pfcrt_30_06_15.fasta, pfcrt_30_06_15.sam, pfcrt_flt_30_06_15.sam, pfcrt_50_30_06_15.fastq) are necessary for producing the CSV file and can therefore be referred to as intermediate files. Note that most of the pipeline's output files are named in the format: *gene_dd_mm_yy*. Where dd, mm, and *yy* are the day, month and year of the current date of analysis, respectively. Typically, the <directory_of_ab1_files> will not contain any intermediate files or output file until the pipeline has ran successfully.

### Results and Discussion

The primary output of this pipeline is the CSV file listing the codons of interest in a specific ADR gene (and their translations) for samples and reference sequences. By successively, analyzing sequencing data for *pfmdr1, pfcrt, dhps*, and *dhfr* the molecular markers of ADR associated with these genes can be detected. The *pfcrt* analysis example, shown above and whose output is in the example_data directory will be used to demonstrate how to work with data produced by this pipeline to determine the frequency of molecular markers of ADR.

Typically, the result for the reference gene is in the last non-empty row of the CSV file. Reporting the codons and codon translations for the reference along with the samples serves as a positive control, to show that the pipeline worked as expected. The reference gene usually contains the wild-type haplotype and can be compared with sample haplotypes to detect variants. Each of these variant haplotypes can be manually verified by visualizing in tablet, as shown in Figure 1. The data in the CSV file can be readily analyzed to determine the frequency of all haplotypes, including those haplotypes that serve as markers of ADR. The R script (adr_snps_analysis.R), in the example_ab1 directory, can be modified to determine these frequencies. Indeed, this script has been used to analyze data in pfcrt_30_06_15.csv (found in the example_data directory). The frequencies of haplotypes in the codon 72 – 76 region of *pfcrt* based on pfcrt_30_06_15.csv are presented in in table 2, below.

**Table 2.** Frequency of *pfcrt* 72–76 haplotypes for 44 samples. Two samples contain mutant haplotypes (CVIET, CVMNT) that contain the K76T mutation, which confers resistance to chloroquine.

| Haplotype | CVIET | CVMNK | CVMNT |
| --- | --- | --- | --- |
| Frequency | 1 | 42 | 1 |

Despite beginning the analysis with 50 samples (50 .ab1 trace files), only 44 of these samples are reported in Table 2. One sample read was too short after quality trimming – its length was less than 19, the minimum remaining sequence length, and was therefore discarded. Additionally, 5 sample reads were filtered out due to poor mapping quality (MAPQ <10). Furthermore, these unmapped sample reads also had low base quality scores, as shown in the plot (Figure 2) produced by FastQC [31].

**Figure 2.**
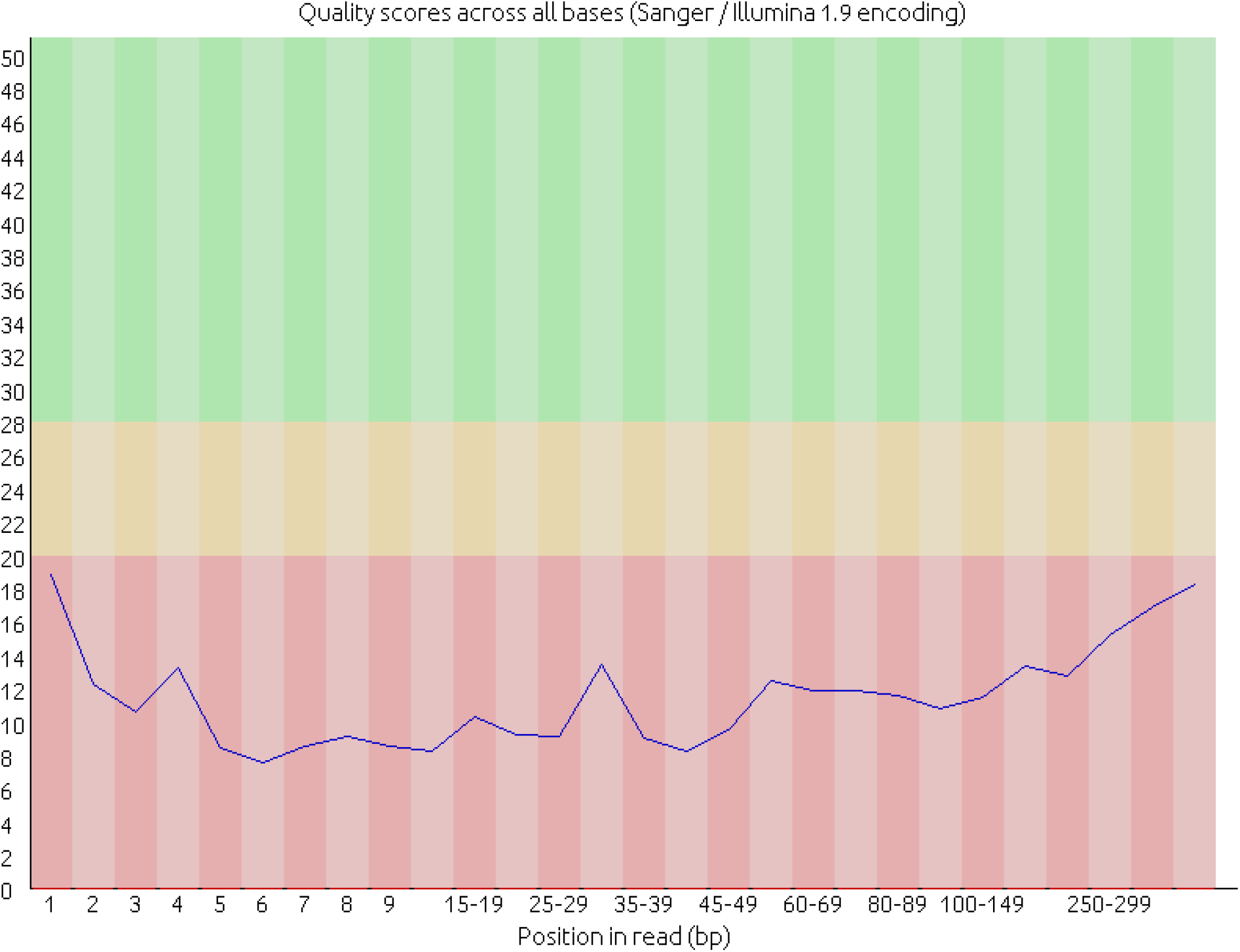
Quality scores per base for unmapped samples. All bases in the unmapped samples have quality score less than Q20.

Manual steps can be taken to characterize samples that ADRes failed to characterize automatically. For instance, unmapped reads can be extracted from the unfiltered SAM file (pfcrt_30_06_15.sam) using the following command, working from the ADRes directory:

./samtools-1.2/samtools view -Sf4 pfcrt_30_06_15.sam > pfcrt_unmapped_30_06_15.sam

This result can then be converted from SAM to FASTQ format using this command [32]:

cat pfcrt_unmapped_30_06_15.sam | grep -v ^@ | awk '{print "@"$1"\n"$10"\n+\n"$11}' > pfcrt_unmapped_30_06_15.fastq

Ultimately, the unmapped reads, in FASTQ format, can be re-aligned to the reference CDS using, bwa bwasw with modified alignment options (executed from the example_ab1 directory):

./bwa-0.7.12/bwa bwasw -z 5 ../pfcrt_pf3d7_cds.fasta pfcrt_unmapped_30_06_15.fastq > bwasw_z5_unmapped_pfcrt_30_06_15.sam

By increasing the value of the z option of bwa bwasw, one of the previously unmapped reads maps to the reference CDS. This sample (41C_CRT_F) was subsequently found to possess the CVMNK wild-type haplotype after converting the SAM file to FASTA format, and parsing the alignment using pfcrt.py:

1. ./sam2fasta.py ./example_data/pfcrt_pf3d7_cds.fasta ./example_data/example_ab1/bwasw_z5_unmapped_pfcrt_30_06_15.sam ./example_data/example_ab1/bwasw_z5_unmapped_pfcrt_30_06_15.fasta
2. ./adr_codons.py -i ./example_data/example_ab1/bwasw_z5_unmapped_pfcrt_30_06_15.fasta -g pfcrt

One way to reduce sample loss due to poor mapping or base quality is to increase sequencing coverage. If samples are sequenced in the forward and reverse directions, the likelihood of losing a sample due to poor quality is reduced. In this example, samples were sequenced in the forward direction only. Therefore, when one read fails a quality test, a sample is lost.

### Conclusions

To detect molecular markers of resistance to common anti-malarials, from Sanger sequencing data, we have developed a functional pipeline comprising widely used tools and customs scripts written in Bash and Python called ADRes. In the near future, the pipeline will be upgraded to support detection of molecular markers to artemisinin resistance based on the kelch13 (K13) – propeller locus [33]. This pipeline has the potential to speed up anti-malarial drug resistance surveillance conducted by research groups with or without specialized bioinformatics staff. The software is open source and is released under a GNU General Public License. Developers seeking to optimize the pipeline for greater functionality and ease of use are welcome to do so.

## Availability and requirements

Project name: ADRes

Project homepage: http://setfelix.github.io/ADRes

Operating system: Linux

Programming languages: Bash 4, Python 2.7

Any restrictions to use by non-academic users: None

## List of abbreviations used

ABI: Applied Biosystems

SNP: Single nucleotide polymorphism

BWA: Burrows-Wheeler Aligner

ADR: Anti-malarial Drug Resistance

CSV: Comma Separated Values

TES: Therapeutic Efficacy Studies

ACT: artemisinin-based combination therapies

AAQs: aminoalcohol-quinolines

SAM: Sequence Alignment/Map

## Authors' contributions

SA wrote the code for the pipeline, and prepared the first draft of the manuscript.

AG conceived the idea of a pipeline, provided example data for analysis, and revised the manuscript.

## Acknowledgements

This work was sponsored by an NIH-NIAID grant 5R01AI099527-02 awarded to AG.

